# Deceptive learning in histopathology

**DOI:** 10.1101/2022.04.21.489110

**Authors:** Sahar Shahamatdar, Daryoush Saeed-Vafa, Drew Linsley, Farah Khalil, Katherine Lovinger, Lester Li, Howard McLeod, Sohini Ramachandran, Thomas Serre

## Abstract

Deep learning holds immense potential for histopathology, automating tasks that are simple for expert pathologists, and revealing novel biology for tasks that were previously considered difficult or impossible to solve by eye alone. However, the extent to which the visual strategies learned by deep learning models in histopathological analysis are trustworthy or not has yet to be systematically analyzed. In this work, we address this problem and discover new limits on the histopathological tasks for which deep learning models learn trustworthy versus deceptive solutions. While tasks that have been extensively studied in the field like tumor detection are reliable and trustworthy, recent advances demonstrating the ability to learn molecular profiling from hematoxylin and eosin (H&E) stained slides do not hold up to closer scrutiny. Our analysis framework represents a new approach in understanding the capabilities of deep learning models, which should be incorporated into the computational pathologists toolkit.

## 1 Introduction

Histopathology image analysis is essential for diagnosing cancer and determining the course of treatment [1]. There is now growing evidence that deep neural networks (DNNs) can at least partially automate this procedure. For instance, DNNs rival expert pathologists in detecting malignant skin lesions [2, 3], diagnosing diabetic retinopathy [4–6], and detecting breast cancer [7, 8]. In each of these cases, DNNs learned to solve straightforward but time-consuming tasks that are already within the expert physician’s repertoire. However, there is now a growing number of reports that DNNs are also capable of solving tasks that are difficult or impossible for pathologists to solve by visual analysis alone. These tasks include profiling the genome of a tumor [9–12] and identifying the originating tissue for cancer of unknown primary [13, 14] (CUP) from tissue morphology. Underlying these findings is the assumption that histopathology images depict patterns of morphological features that are impossible for experts to detect by eye, but which DNNs have the capacity to learn to identify. This assumption has not been systematically tested, raising the possibility that DNNs solve histopathological tasks by exploiting morphology that is indirectly or spuriously related to disease.

The trustworthiness of DNNs is an open problem in computer vision because DNNs often exploit idiosyncratic or spurious correlations between images and labels to solve visual tasks [15, 16], a strategy which can lead to high accuracy on a single benchmark dataset but fails to hold up under more rigorous testing. Such a reliance on spurious features has caused many notable errors in estimating the abilities of animals and machines. For example, the notable case of “Clever Hans” [17], the horse who relied on nonverbal cues from his trainer to solve simple arithmetic, has been evoked as an analogue for the proposensity of the learned strategies of DNNs to be misaligned with humans [16]. To avoid such errors when developing DNNs for clinical applications, it is standard practice to validate models by visualizing the morphological features that drive their decisions [9, 11, 12, 18–22]. If a model relies on similar morphological features as an expert pathologist for a single well-defined task, the visual strategy it has learned for histopathological analysis is aligned with humans and trustworthy. While this approach is effective for comparing the strategies of humans and machines on established histopathological analysis tasks like tumor detection, it is poorly suited for tasks that are difficult or impossible for expert pathologists, which do not have established morphological criteria. Indeed, for difficult tasks like molecular profiling from hematoxylin and eosin (H&E) slides, the features DNNs use to render their decisions are often presented without any biological ground-truth for comparison [9, 11, 12, 18–22], possibly because the appropriate biological data is impossible or very costly to acquire. Here, we challenge this standard for assessing DNN visual strategies in histopathological image analysis to identify tasks where they are trustworthy or deceptive. If a DNN is trustworthy, its visual strategy will closely align with expert pathologists on standard tasks, or reflect novel disease-relevant features on tasks that are difficult or impossible for expert pathologists. On the other hand, a deceptive DNN will rely on visual features that are unrelated to the underlying biology and ultimately useless for the clinic or research.

We systematically test the trustworthiness of DNNs by measuring the visual strategies they learn after being trained to solve multiple histopathology analyses ranging in difficulty, from straightforward tumor detection to tasks that are potentially impossible for expert pathologists: determining the molecular profile and primary tissue of a tumor (CUP). Next, we develop a standardized evaluation framework to distinguish between trustworthy and deceptive visual strategies of DNNs trained to automate these analyses. Our general approach is to infer the tissue morphology driving DNN decisions, then compare this morphology to gold-standard diagnostic criteria for the same task. For molecular profiling, where there is no such gold standard — common wisdom is that expert pathologists cannot do this from tissue morphology alone — we use laser capture microdisection (LCM [23]) to collect ROI-based genomic labelings on adjacent WSIs to ones for which we have full-slide genomic labels. For determining the primary tissue of a tumor, we restrict our analysis to tumor subtypes that contain no tissue-specific features. Our approach fills a void in the rapidly growing field of deep-learning-based histopathological image analysis: revealing tasks that are vulnerable to deceptive learning and as of now unsuitable for automation with state-of-the-art deep learning models and training routines.

## 2 Results

### Overview

We investigated the trustworthiness of DNNs trained for histopathological analysis by developing a standardized evaluation framework, which consists of two steps. **Step 1:** models are tested within and outside of the training distribution to identify trivial shortcut strategies by measuring generalization performance. **Step 2:** the morphological features each model relies on for solving a task are inferred, validated for their importance to a model’s decisions, then compared to the task’s gold-standard diagnostic criteria to detect deceptive visual strategies.

We scrutinize four popular DNNs trained to solve multiple image analysis tasks posed on a novel dataset of 221 histology WSIs. These WSIs depict H&E stained formalin-fixed paraffin-embedded (FFPE) tissue from 182 patients diagnosed with lung adenocarcinoma at the Moffitt Cancer Center (Methods). We refer to this dataset as the Moffitt dataset. Our DNNs were 18, 34, and 50-layer versions of the ResNet V2 [24] and the ShuffleNet V2 [9, 11, 24].

Each of the DNNs was trained to solve three analysis tasks posed on lung adenocarcinoma slides that range in difficulty for expert pathologists, from potentially impossible to a standard histopathological analysis: (i) molecular profiling of KRAS versus EGFR mutated tumor (KRAS-mutation), (ii) determining the primary tissue of a tumor (CUP), and (iii) classifying lung adenocarcinoma versus benign tissue (tumor detection, task denoted as tumor-normal). As detailed in the results, we utilized a combination of molecular and expert pathologist annotations in order to pose these three distinct tasks on the same slides. For each task, we analyzed the performance and interpretability of the four DNN architectures on WSIs from our novel “Moffitt dataset,” which was held out from training or model selection (Figure 1). To measure out-of-distribution generalization, we tested models on the public TCGA slide image dataset (173 total slides; https://www.cancer.gov/tcga), which includes clinical, genomic, and histologic data for lung adenocarcinoma.

**Fig. 1 Figure 1.**
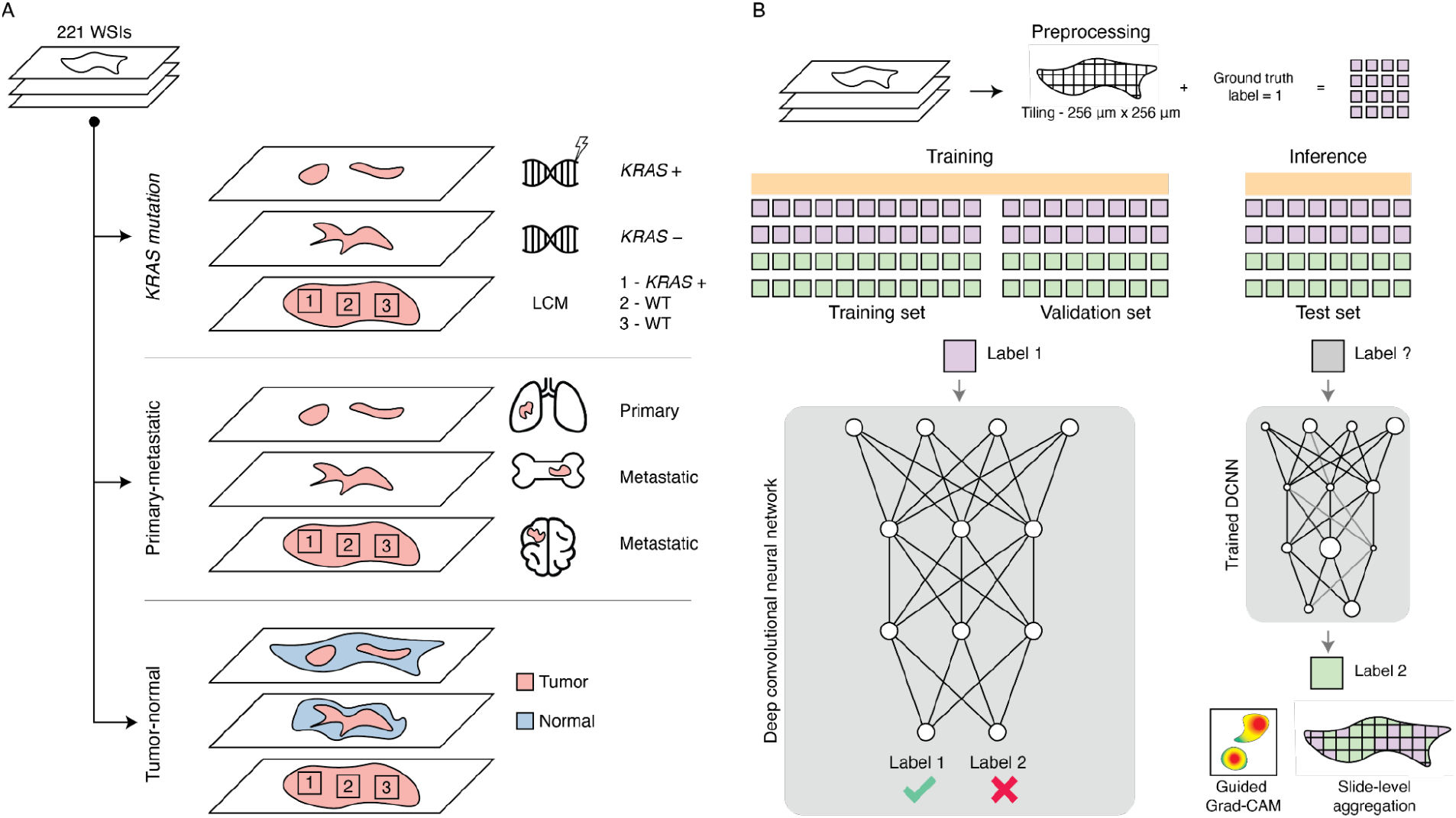
Overview of classification tasks and deep learning framework. (**A**) Schematic showing the three classification tasks. For the KRAS-mutation task data, we divided our dataset into WSIs from tissue that had mutations in the KRAS gene (KRAS+) and WSIs from tissue that did not have mutations in the KRAS gene (KRAS-). We collected additional sequencing information using laser capture microdissection (LCM) to dissect and sequence arbitrarily selected regions of interest (depicted by boxes 1-3) for mutations in KRAS. For the primary-metastatic task data, we identified WSIs that were biopsied from the lung (primary) or from other sites but with a known pulmonary origin (metastatic). Only tissue inside the tumor annotated area was used in this task. Lastly, for the tumornormal task data, we identified tumor and normal tissue in all whole slide images (WSIs) using pathologist tumor annotations. Tumor tissue is colored pink, and normal tissue is colored blue. (**B**) All WSIs were tiled into non-overlapping 256 *μm* × 256 um tile images, corresponding to 512-pixel × 512-pixel image patches at 20× magnification. Each image patch was assigned a different ground-truth label for each task. The image patches were divided into training, validation, and test folds; each patient’s image patches belonged to a single fold. The image patches in the training and validation sets were used to train and select hyperparameters, whereas image patches from the test fold were used for model evaluation.

Each WSI in the Moffitt dataset and TCGA yielded, on average, about 2,000 non-overlapping 512-pixel × 512-pixel image patches. We partitioned WSIs from the Moffitt dataset into separate cross-validation folds for training, model selection (validation), and testing (each fold contained unique WSIs, see Methods). Models were then trained and evaluated on image patches extracted from these WSIs, as is standard practice in the field due to the computational complexity of training directly on WSIs [9, 11, 12, 18–22].

### Task 1: DNNs trained for molecular profiling are vulnerable to deceptive learning

We began by analyzing the performance of DNNs trained to do molecular profiling on H&E WSIs. Molecular profiling is an important step for determining the course of treatment for the disease which requires expensive and time-consuming genomic panels. Recent work has shown that genomic mutations can be reliably decoded from morphology by DNNs [9, 11, 12]. Are these DNNs and their putative successes trustworthy or deceptive?

We modeled clinically significant genes to remain consistent with prior work on the task [9, 11, 12], focusing specifically on whether or not DNNs could discriminate mutations of genes that do not co-occur in the Moffitt dataset or TCGA: KRAS and EGFR. We trained models for binary classification, where KRAS mutations were the positive class, and EGFR mutations were the negative class. All image patches for training and evaluation were taken from inside tumor-annotated regions of WSIs. Models were evaluated by recording area under the receiver operator characteristic curve (ROC-AUC) and class-weighted accuracy.

To evaluate DNNs on molecular profiling, we first measured their performance within and outside of the training distribution. All DNNs were significantly above chance at detecting KRAS mutations at the slide-level in the Moffitt dataset (*p* < 0.05 over 1000 bootstrap replicates, where chance = 0.5 for both weighted accuracy and ROC-AUC; Table 1). There were no significant differences between the performance of the models, and they all rendered similar decisions on the task, indicating that model architecture differences did not translate into qualitatively different task strategies (all model-to-model Pearson correlations exceeded *ρ* = 0.68, *p* < 0.001, SI Table 1). The models also performed significantly above chance when tested out-of-distribution on the TCGA (SI Table 2). In other words, all of the DNNs we tested learned similarly generalizable strategies for detecting KRAS at the WSI level. These results are consistent with multiple other applications of DNNs for molecular profiling10–12, which indicate that DNNs are significantly above chance in profiling WSI genomes, while still far below the error rate of standard molecular tests (most assays can detect somatic mutations at greater than 95% sensitivity and specificity [25]).

**Table 1.** DNNs successfully detect KRAS mutations at the whole-slide level, but fail to localize KRAS mutations within a slide. Slide predictions are calculated from the median of logit scores across all image patches in a WSI. LCM-captured image patches are evaluated independently. For each metric, we report the 95% confidence interval using 1000 bootstrap replicates. Statistical testing against chance accuracy (0.5) is denoted by asterisks: ** = *p* < 0.01, * = *p* < 0.05.

| Analysis | Score | ResNet-18 | ResNet-34 | ResNet-50 | ShuffleNet |
| --- | --- | --- | --- | --- | --- |
| WSI (whole-slide level) | Weighted accuracy | 0.59 [0.52 - 0.68]* | 0.60 [0.53 - 0.68]* | 0.60 [0.53 - 0.69]* | 0.63 [0.54 - 0.71]* |
|  | ROC-AUC | 0.69 [0.59 - 0.78]** | 0.66 [0.55 - 0.76]* | 0.70 [0.60 - 0.78]** | 0.67 [0.57 - 0.77]** |
| LCM Patch (localization) | Weighted accuracy | 0.51 [0.45 - 0.57] | 0.38 [0.33 - 0.44] | 0.45 [0.40 - 0.50] | 0.47 [0.41 - 0.53] |
|  | ROC-AUC | 0.50 [0.43 - 0.57] | 0.40 [0.33 - 0.47] | 0.44 [0.37 - 0.50] | 0.46 [0.39 - 0.53] |

The above-chance generalization performance of the DNNs means that it is unlikely that they learned to detect KRAS through a trivial shortcut [15], such as experimental batch effects or systematic variations in how the slides of different patients were handled and prepared. However, it is still possible that the DNNs learned to rely on a deceptive visual strategy, exploiting visual features that are correlated to KRAS mutations but not related to the underlying biology. Because there are no gold-standard morphological phenotypes for different mutations, it is difficult to detect such deceptive learning on this task.

To investigate whether our DNNs learned deceptive visual strategies for detecting KRAS mutations, we gathered additional sequencing data on 20 patient WSIs. We collected 10-20 1×1mm regions-of-interest (ROIs; n=216) at positions distributed within the tumor of each of these WSIs using LCM. Next, we sequenced each ROI for KRAS and EGFR mutations (see Methods), providing us an estimate of the spatial distribution of these mutations in the WSIs. There were 90 patches with KRAS mutations, 109 patches with EGFR mutations, and 17 wild-type (WT) patches without mutations in either gene. KRAS mutations were spatially heterogeneous, and four of the slides that were labeled as KRAS mutation based on their whole-slide panel contained regions of wild type (17 total patches).

No DNN exceeded chance-level on the LCM patches, and all performed significantly worse than on WSIs (Table 1 and Figure 2). These results raised the question: what morphology had the DNNs learned to rely on to perform above chance at the level of WSIs?

**Fig. 2 Figure 2.**
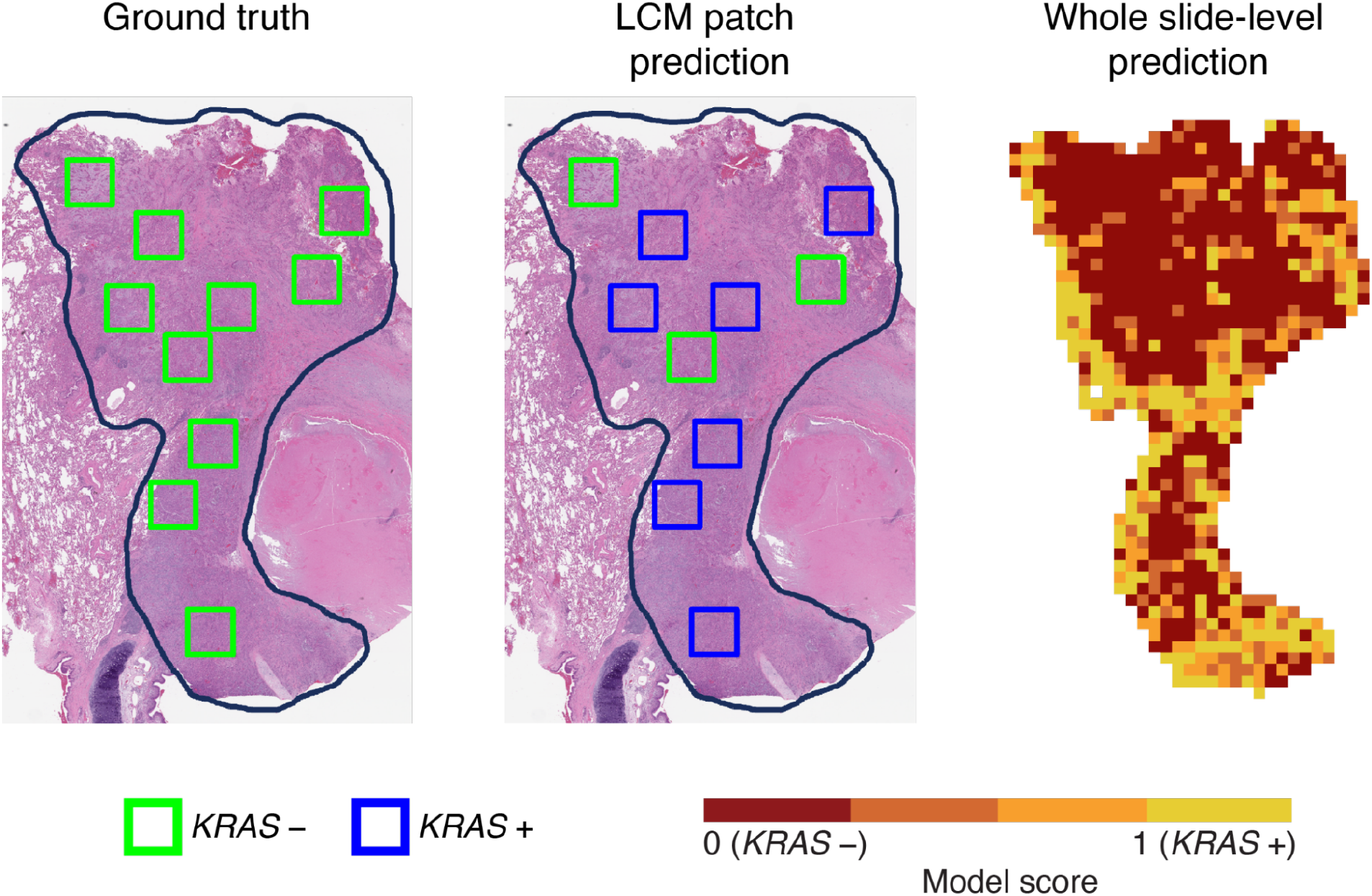
DNN KRAS mutation localization is inconsistent with LCM sequencing. (Left and middle) H&E stained whole-slide images with tumor outlined in black, and boxes representing locations of LCM, which are colored according to their molecular labels. The left panel depicts the ground truth labels, and the middle the predictions of the ShuffleNet model, which accurately classifies three LCM patches as KRAS negative (green) and incorrectly classifies the rest of the seven patches as KRAS positive (blue). On the right, the heatmap is colored according to the ShuffleNet model score; a model score of 0 (maroon squares) corresponds to no KRAS mutation detected and 1 (yellow squares) corresponds to KRAS mutation detected. Areas outside of the annotated tumor or where there were no cells are white. Despite the heterogeneity of the model’s predictions at the LCM level, the model’s whole slide decision correctly identifies this as a KRAS wild-type tumor.

A popular approach to interpret DNN decisions is to compute the gradient of a model’s output (*z*) with respect to the input image (*x*): 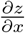. This vector of gradients captures the importance, or “saliency” of every pixel in the input image to the model’s decision for that image, where larger (absolute) gradient values denote more importance [26–29]. Model saliency maps are noisy estimates of feature importance, and subsequent advancements in explainable artificial intelligence have involved modifications of the gradient to reduce noise in these maps. A particularly effective approach for biomedical imaging [30] is Guided Gradient-weighted Class Activation Mapping (GradCAM [31]), which we use here to identify morphological features that drove DNN molecular profiling decisions. We specifically focus on interpreting Shufflenet’s decision making, because it performed best of the models we tested at detecting KRAS mutations on WSIs.

We applied GradCAM to the predictions of Shufflenet on WSIs in the Moffitt test set. In order to identify the types of features the model relied on for its decisions, we also passed these slides through HoVer-Net [32], a DNN trained to segment nuclei in H&E images. Nuclear features in H&E slides are commonly used to assess a host of different diseases [33]. By measuring the overlap between GradCAM maps for KRAS detection and nuclear features, we reasoned that it would be possible to measure the extent to which model decisions relied on each type of feature. On average, only 14% of model GradCAM pixels for KRAS detection overlapped with tumor nuclei predictions (this metric ranged from 11-15% for the three other models).

We also explored the correlation between model decisions and histological subtype, a morphological feature that is known to be weakly associated with KRAS mutations [21]. Histological subtyping, unlike molecular profiling, is straightforward for expert pathologists and is detectable by DNNs [34]. It is possible that DNNs trained for KRAS detection could not find strong morphological signals for KRAS, and instead learned to rely on weakly correlated subtypes. If this is the case, the drop in DNN performance when tested on WSIs versus LCMs could be explained by the correlation strength — or lack thereof — between subtypes and mutations on WSIs versus LCMs. To test for this possibility, we annotated the histological subtypes present in each LCM patch image and the WSIs they came from. We found six total subtypes: acinar, lepidic, solid, papillary, micropapillary, and mucinous. We next computed the correlations between these subtypes and the presence of KRAS mutated vs. KRAS wild-type in the LCMs and WSIs.

We used logistic models to regress the presence or absence of a KRAS mutation in each WSI onto its annotated histological subtypes. Consistent with prior work [21], we found a significant (one-tailed) association between the micropapillary subtype and KRAS wild-type (*z* = −1.675, *p* = 0.044), and a significant correlation between the solid subtype and KRAS mutations (*z* = 2.026, *p* = 0.026). However, after repeating this analysis on the LCM patch images, we found no significant correlations between histological subtype and KRAS mutated or KRAS wild-type. This means that a model which has learned to detect KRAS mutations by focusing on subtype at the whole slide level should fail to generalize to LCM patches, where these correlations are absent. We verified that the Shufflenet adopted this strategy by fitting a linear model to regress the WSI histological subtypes onto its KRAS mutation predictions (in logits). The model’s predictions were significantly correlated with the solid subtype (*z* = 1.814, *p* = 0.035) despite being trained to detect KRAS mutations, validating our hypothesis that these models tend to rely on histological subtype to detect mutations.

DNNs are significantly above chance at detecting mutations from histology in WSIs because they learn to classify histological subtypes that are correlated with those mutations. Subtypes are straightforward to classify by eye for expert pathologists and are only weakly associated with genetic mutations. As we demonstrate, these correlations exist at the whole-slide level but disappear at granular levels of analysis. That DNNs learn to rely on subtypes as a shortcut for molecular profiling means that their predictions of morphology related to genetic mutation are deceptive. This is especially an issue when models are asked to render decisions at a different level of analysis than they were trained, such as predicting mutations for particular cells after being trained on whole slides.

### Task 2: DNNs can reliably identify the primary tissue of a tumor

Are there other tasks that are difficult or impossible for expert pathologists that DNNs can learn to reliably solve? To address this question, we next asked how effectively DNNs can classify the primary tissue of a tumor. Cancer of unknown primary (CUP) describes tumors for which the primary anatomical site cannot be determined. Because this information is critical for modern therapeutics, clinicians often turn to expensive and time-consuming genetic or transcriptomics analyses. However, it has recently been suggested that DNNs can diagnose the origin of the primary tumor from morphology alone [13] even though expert pathologists are incapable of doing this task by eye.

We tested whether the ability of DNNs to detect the primary tumor origin is trustworthy or not by posing a version of the task on the same lung adenocarcinoma slides we used to investigate molecular profiling. We first split up the slides into primary and metastatic lung adenocarcinoma. Next, we restricted our set of slides for model training to only contain those which had tumors composed of the solid histological subtype, which controls against the presence of trivial features for detecting tissue type. We extracted image patches from tumors in these slides for model training. Finally, we developed an additional test set of slides containing image patches only from sheets of tumor cells in the solid subtype tumor, which is the strictest possible criterion for controlling tissue-specific shortcuts for solving the task.

As with the molecular profiling task, we first measured the generalization performance of DNNs trained to solve this task. All of the DNNs performed significantly above chance in differentiating primary and metastatic lung adenocarcinoma in the Moffitt dataset, reaching between 60% and 72% accuracy (Table 2), and rendering similar decisions to each other: the Pearson correlation coefficient between all models’ decisions ranged between 0.74 and 0.77 (*p* < 0.001 for all model pairs according to randomization tests [35]).

**Table 2.** All four DNNs can differentiate primary lung adenocarcinoma images from metastatic adenocarcinoma images. For each metric, we report the 95% confidence interval using 1000 bootstrap replicates. Statistical testing against chance accuracy (0.5) is denoted by asterisks: ** = *p* < 0.01.

| Analysis | Score | ResNet18 | ResNet34 | ResNet50 | ShuffleNet |
| --- | --- | --- | --- | --- | --- |
| Tile | Weighted accuracy | 0.60 [0.60-0.61]** | 0.72 [0.71-0.72]** | 0.69 [0.68-0.69]** | 0.71 [0.71-0.72]** |
|  | ROC-AUC | 0.62 [0.61-0.62]** | 0.82 [0.81-0.82]** | 0.74 [0.73-0.74]** | 0.73 [0.72-0.73]** |

We next evaluated out-of-distribution generalization of these DNNs on the TCGA dataset. All subtypes were included as we did not have histological subtype annotations for many of the WSIs in the TCGA dataset. In addition, since the TCGA only contains primary lung adenocarcinoma, we measured performance by recording true positive rates. All models maintained true positive rates that were significantly greater than chance in this generalization task (*p* < 0.05), with tile true positive rates ranging from 70% to 84%.

We focus our analyses here on the best performing ResNet34 (see SI for the other models) and adopted the same evaluation strategy from the molecular profiling task to identify and analyze morphological features that the DNNs use for differentiating primary from metastatic lung adenocarcinoma. However, while it is apparent that the model learned to target tumor cells, the differences between the features selected in its GradCAM maps for primary and metastatic tissue was small, with less fibrotic tissue in the metastatic versus primary images (Figure 3).

**Fig. 3 Figure 3.**
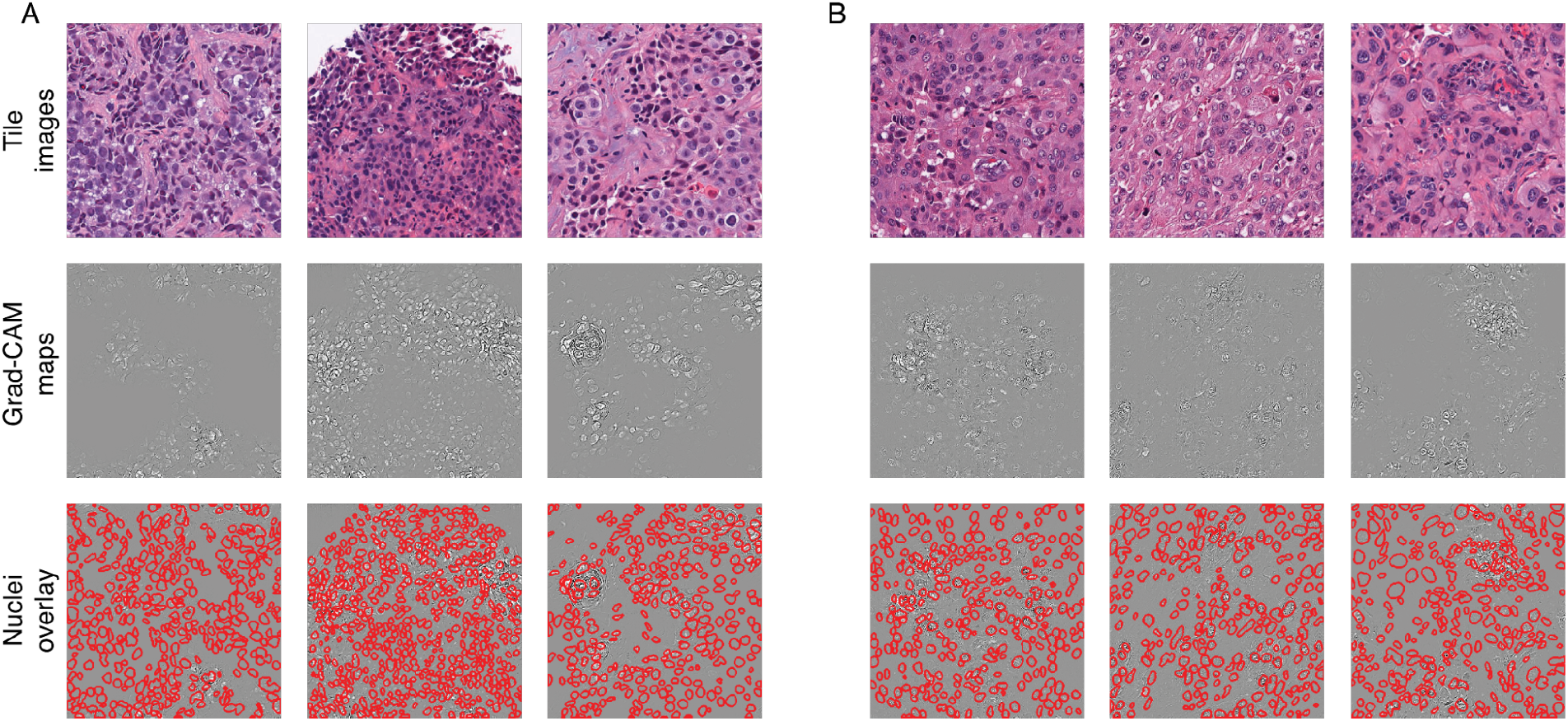
Deep learning models use nuclear and non-nuclear features to identify the primary tissue of a tumor. The first row shows three representative tile images from (A) metastatic and (B) primary tile images (WSIs). Guided GradCAM maps from ResNet34 for each tile image are shown in the second row. The last row depicts GradCAM maps with segmented nuclei outlined in red..

In the absence of obvious morphological criteria, we next tested whether nuclear features were driving model decisions. We found that model performance was significantly reduced when we masked DNN activities corresponding to the locations of nuclei segmented by HoVer-Net (unmasked weighted accuracy 71.3% vs. nuclei masked weighted accuracy 63.9%, *p* < 0.05) but the performance remained significantly above chance. These results suggest that the model learned a visual strategy for classifying the primary tissue of a tumor which relied on both nuclear and non-nuclear features.

To summarize, we found that DNNs can identify a tumor’s primary tissue by leveraging a strategy that is at least partially trustworthy. When tested on sheets of solid subtype tumor cells, models were surprisingly significantly above chance, utilizing a combination of nuclear and non-nuclear features to achieve this performance. Our findings on this task indicate a more bullish outlook for DNNs trained to identify a tumor’s primary tissue than those trained for molecular profiling (Task 1). Our models predict that there are reliable morphological features for identifying a tumor’s primary tissue even though the task is exceedingly difficult or impossible for expert pathologists.

### Task 3: DNNs learn a robust strategy to detect tumors

Finally, we turned to a well-studied task in computational pathology: detecting lung adenocarcinoma in WSIs. This task is routine for expert pathologists, and DNNs trained to solve it have rivaled the performance of experts [9, 36–38]. To test whether or not DNNs adopt deceptive visual strategies to solve tumor detection, we posed the task on the same datasets we used for molecular profiling and classifying primary tissue.

We trained and evaluated models on image patches extracted from WSIs, as was done in prior work [9, 36–38]. After training and evaluating on the Moffit dataset, we found that the ShuffleNet and all three ResNets performed similarly to the state of the art [9, 36–38], reaching approximately 88% weighted accuracy and 0.95 AUROC (Table S4). We focused subsequent analyses on the best-performing model, the ResNet18.

When evaluated out-of-distribution on lung adenocarcinoma WSIs in TCGA, the ResNet18 once again performed well, reaching nearly 82% accuracy and 0.90 AUROC (Table S4). This decrease in performance is relatively small but statistically significant (95% confidence intervals do not overlap, *p* < 0.05), and likely reflects experimental measurement differences between the two datasets, such as their use of different staining protocols [39]. Notably, however, the model’s out-of-distribution generalization is still significantly better than chance (*p* < 0.001). As in the other two tasks, all models trained for tumor detection rendered similar decisions to each other on the Moffitt dataset and out-of-distribution generalization on the TCGA dataset. The Pearson correlation coefficient between models’ decisions on every WSI was between 0.79 and 0.89 for both datasets (*p* < 0.001 for every pair).

The ResNet18 accurately localized tumor tissue in Moffitt and TCGA WSIs (Figure 4). Image patches from the Moffitt dataset, for which ResNet18 is most confident contain tumor tissue, also have morphological features associated with lung adenocarcinoma39, such as irregular and enlarged nuclei, prominent nucleoli, and high nuclear to cytoplasmic ratio (Figure 5A). Moffitt image patches which ResNet18 is most confident are benign, depict benign alveolar septa (the tissue that separates the alveoli, or air sacs) with empty air space and maintained structure, lined by benign type I (flat cells) and type II (cuboid) epithelium (Figure 5C). Overall, the distinguishing features in the tile images with high and low model scores correspond to features that pathologists use to distinguish between malignant and benign tissue [38].

**Fig. 4 Figure 4:**
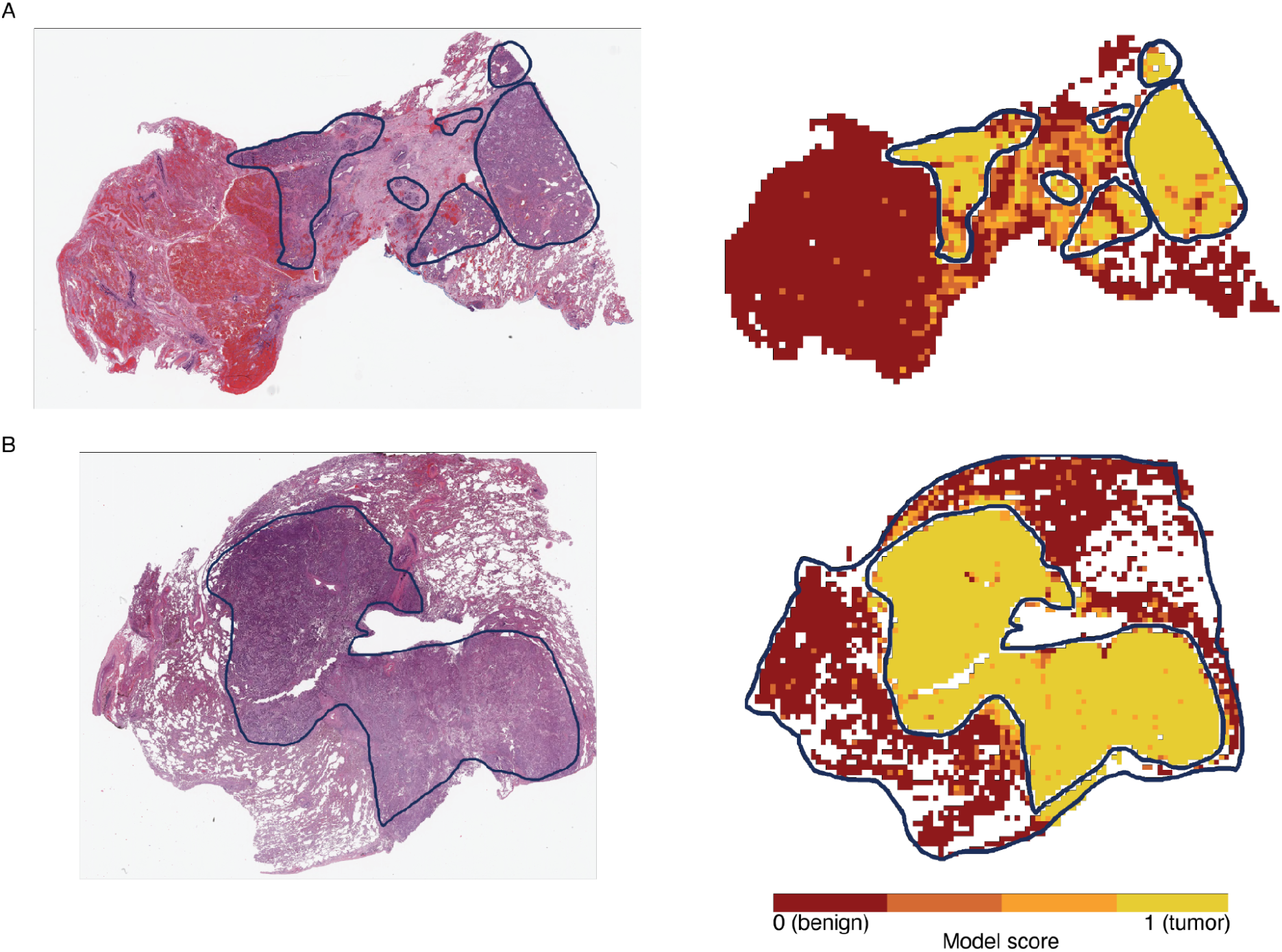
Prediction heatmaps for ResNet18 trained to detect lung adenocarcinoma. Two whole slide images (WSIs) of lung adenocarcinoma are annotated for tumor regions in blue. The heatmaps in the right column are colored according to the ResNet18 model score. A model score of 0 corresponds to benign tissue and 1 corresponds to tumor tissue.

**Fig. 5 Figure 5:**
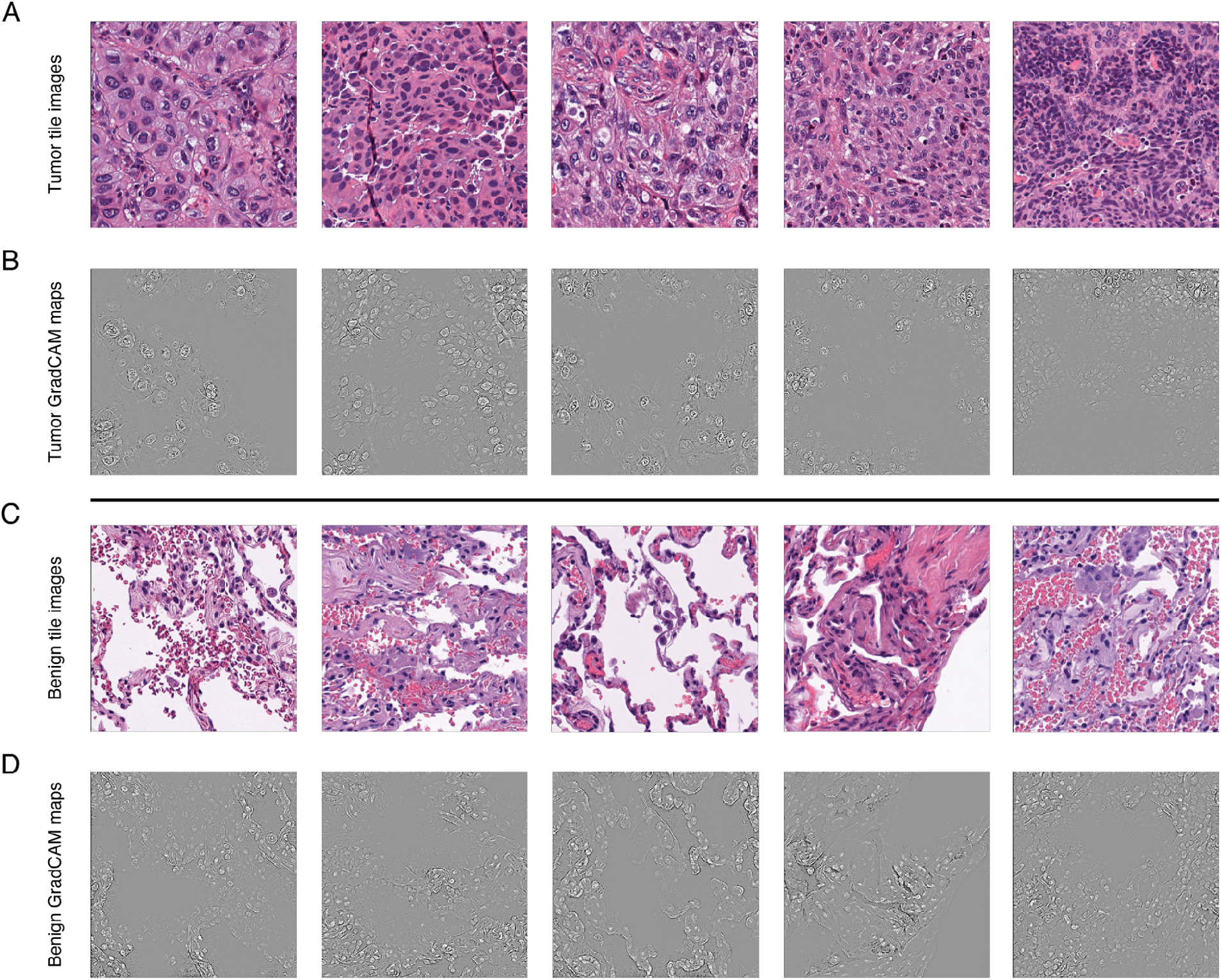
Deep learning models use nuclei to differentiate between tumor and benign tile images. **A.** Representative tile images inside the tumor-annotated area with high model scores (higher model score = more tumor-like). All model scores are greater than 0.999. **B.** Tumor class GradCAM maps for each of the tile images in panel A. The GradCAM signals often overlap with cell nuclei, and more specifically tumor cell nuclei. **C.** Representative tile images outside the tumor-annotated area with low model scores (lower model score = more benign-like). All model scores are less than 1e-3. **D.** Benign class GradCAM maps for each of the tile images in panel C. The GradCAM signals often overlap with cell nuclei and outline alveolar septal tissue.

To establish whether nuclear features were driving model decisions, we again masked DNN activities corresponding to the locations of nuclei in image patches. Ablating all tumor nuclei significantly reduced weighted accuracy from 88.3% to 72.4%. We repeated this experiment by permuting the tumor nuclei masks; the accuracy dropped to 85.7%. Ablating all nuclei reduced weighted accuracy to chance-level at 49.8%, while randomly permuting the masks reduced accuracy to 59.2%.

Lung adenocarcinoma classification with DNNs is trustworthy. All DNNs we tested perform well both within and outside of the training distribution, and render similar decisions (model-to-model correlations are all between 0.79 and 0.89, p j 0.001). These decisions are driven by, and depend on, nuclear morphology that is consistent with textbook and gold-standard criteria for detecting lung adenocarcinoma in histopathology. One possible explanation for the consistent and strong performance of DNNs on lung adenocarcinoma classification is that annotations are at the pixel level, rather than at the whole-slide level like in molecular profiling or classifying the primary tissue of a tumor. This level of granularity in annotations on training data has been found to drive DNNs towards consistent and generalizable visual strategies in object classification tasks29 and may be necessary to avoid deceptive learning.

## 3 Discussion

There is growing consensus that DNNs can automate tasks on biomedical imaging data, achieving performance that matches or exceeds human experts [30, 40]. It is also becoming clear that these DNN achievements are due to visual strategies that are not necessarily aligned with those used by human experts [26, 28, 30]. When humans and machines use different strategies to solve tasks it can be a positive development, with the chance to reveal new insights into biology, and generate testable hypotheses for understanding the development of disease31. But there is no guarantee that the visual features that machines but not humans use to solve tasks are meaningful. There has been extensive work demonstrating that DNNs are in fact vulnerable to learning shortcuts in standard computer vision tasks posed on natural images, like object recognition [15, 41]. These shortcuts represent visual strategies that achieve high performance by focusing on biases that are unique to the training and testing data — from low-level cues like lightness, contrast, or color to object-centric cues like their size in pixels or common appearance in a specific context. However, little is known about DNN shortcut learning in the context of biomedical imaging, and specifically, histopathological analysis. Through our experiments, we have begun to address this critical question of shortcut learning in histopathological analysis, and demonstrated the steps needed to avoid it. In contrast to proposals from computer vision that shortcut learning can be avoided by simply testing model generalization out-of-distribution [15], we find that it is also essential to analyze the morphological features used by models to solve tasks to ensure that their performance is not exploiting a biologically trivial and ultimately deceptive visual strategy.

There is now evidence that DNNs can solve histopathological tasks that expert pathologists cannot. One foundational example of this trend was the recent demonstration that DNNs can automatically learn molecular profiling from morphology [9, 11, 12]. These models could potentially revolutionize oncology, providing rapid accurate prognoses that replace the expensive and time-consuming molecular panels that are standard in the field, and opening up new vistas for studying cell-cell interactions in the development of cancers. However, upon further examination of these findings, we see evidence that DNNs learn molecular profiling by focusing on a shortcut rather than a previously unknown morphological pattern. DNNs render molecular decisions by categorizing the morphological subtype of tumors, which has a weak association with genomic mutations that has been known for at least a decade [21]. DNNs learn to rely on this shortcut because they are trained to associate all of the potentially genetically heterogeneous image patches from a single WSI with a single sequence taken from the entire WSI. When mutation predictions from these models are tested at a more granular level than the WSI, such as the LCM image patches we introduce in this work, the correlation between subtype and mutation is weakened which causes model performance to drop to chance. DNNs learning to focus on subtype rather than a stronger morphological signal for molecular profiling is not necessarily a shortcut — subtype is not completely spurious — but understanding this visual strategy puts a low ceiling on the utility of DNNs for molecular subtyping using existing datasets and training routines. For this reason, we refer to this DNN strategy as deceptive, and significant work is needed to develop trustworthy DNNs for extracting genomic insights from morphology. We release all the images, labels, and LCMs from our Moffitt Dataset to support this goal.

Not all tasks that are difficult or impossible for expert pathologists are vulnerable to shortcut learning in DNNs. When training DNNs to identify the originating tissue for cancer of unknown primary [13, 14] (CUP) on the same WSIs we used for molecular profiling, we found that models learned a robust visual strategy that did not exploit shortcuts. The models learned a novel combination of nuclear and non-nuclear features to solve the task, which generalized effectively, and which will need additional experimental study to understand their relationship to the development of lung adenocarcinoma. Overall, these findings indicate that recent findings on the ability of DNNs to learn to identify the originating tissue of CUP [13, 14] are trustworthy and an important line of future computational research.

Our success on tumor detection, while not novel, points to a general strategy for ensuring that DNNs are trustworthy in histopathological analysis. In that task, unlike molecular profiling or CUP, annotations are provided at the level of pixels in WSIs. Such a “per-pixel” labeling strategy has proven successful in training DNNs for segmentation tasks on natural images, and can promote visual strategies that align with human perception [28, 42]. When this level of extensive model supervision is not available, because such data is expensive or difficult to gather, the explainability framework we laid out in this paper is necessary for distinguishing between trustworthy and deceptive DNN visual strategies.

Our work shows the utility and dangers of applying deep learning models to histopathology tasks. When training DNNs for histopathological analysis, practitioners should have access to high-quality labeled data as well as explainable artificial intelligence methods to build robust and clinically useful models. Our mixed findings on the trustworthiness of recent efforts in histopathological analysis underscores this message, and indicates that the field needs a far greater emphasis on model interpretability to create automated systems that can aid biomedical research and increase the efficiency of expert pathologists in the clinic.

## 4 Methods

### Patient cohorts

#### Moffitt cohort

The Moffitt cohort includes patients at Moffitt Cancer Center (MCC) diagnosed with adenocarcinoma of pulmonary origin that also have associated molecular profile data from 01/01/2011 to 06/09/202044. Participants were included if they satisfied the following criteria. (*i*) The patient must have a pathologic diagnosis of adenocarcinoma with a pulmonary origin. The adenocarcinoma may involve any organ as long as pathology reports it as primary lung adenocarcinoma (e.g., a brain tumor that is pathologically confirmed as metastatic lung adenocarcinoma). (*ii*) The aforementioned adenocarcinoma must have had any sort of molecular profiling demonstrating either the presence or absence of KRAS mutations or EGFR mutations. (iii) A tissue glass slide associated with the formalin-fixed paraffin-embedded (FFPE) tissue block sent for molecular testing must readily be available through the MCC Pathology Department. Participants were excluded from the cohort if they satisfied any of the following criteria: (i) they had a concurrent pathologic diagnosis of another cancer, (ii) no associated molecular data, (iii) or mutations in HRAS and NRAS genes.

Slides that passed the inclusion/exclusion criteria were reviewed for quality and scanned at 20 × via the Aperio AT2 high-volume digital whole slide scanner. This digitization process produced a digital SVS file per slide scanned. These SVS files are referred to as whole slide images (WSIs). The WSIs were annotated for tumor regions of interest areas using a virtual pen via the Aperio ImageScope pathology slide viewing software. The final annotations were stored in XML files.

#### TCGA cohort

Diagnostic slides for all TCGA lung adenocarcinoma patients were downloaded from the GDC (https://gdc.cancer.gov) in SVS format. Seven patients were excluded secondary to an unacceptable prior treatment, an item not meeting the study protocol. Nine SVS files without magnification data were excluded. Ten SVS files with extensive pen marks were excluded.

### Molecular labels

#### Moffitt cohort

The associated molecular profiling for each WSI was obtained for each patient through requests made to Moffitt’s Collaborative Data Services (CDS). The data came from four different sequencing strategies: FoundationOne CDx assay (Foundation Medicine, Cambridge, MA, USA), Moffitt STAR, TruSight Tumor 15 (Illumina, USA), and in-house pyrosequencing [43]. Only mutations that were labeled as clinically significant by each respective kit were included in the final dataset.

#### TCGA cohort

TCGA molecular alteration labels were derived from the public mutation annotation file prepared by [44]. All intronic and silent mutations were excluded. Patients with mutations in EGFR as well as in a RAS family gene (KRAS, NRAS, HRAS) were excluded. To match our Moffitt cohort, we excluded patients without clinically significant mutations in either EGFR or KRAS (as determined by COSMIC annotations).

### Laser capture microdissection

Twenty-one slides with minimal histological artifacts and abundant tissue were chosen (11 with KRAS mutations and 10 EGFR mutations). For each slide, 10 to 20 1mm × 1mm regions of interest (ROI) in the tumor-annotated area were further annotated. The ROIs were spatially distributed throughout each slide at randomly selected locations within the tumor.

Unstained FFPE tissue sections, prepared on polyethylene napthalate membrane slides, were depariffinized and dehydrated by dipping in 100% xylene for two minutes. Slides were allowed to air dry for five minutes. Selected regions of interest (ROIs) were micro-dissected from the tissue using an Acturus XT Laser Capture Microdissection (LCM) system (Life Technologies Corp., Carlsbad, California) with UV laser cut only. The UV laser precisely cut the PEN membrane around the ROI and the micro-dissected tissue was transferred to a 0.5ml microfuge tube. All ROIs micro-dissected in the study were 1 mm^2^ in size and matched to ROIs annotated by the study pathologist on WSIs representing a sequential section of the tissue. Each ROI was sequenced for KRAS and EGFR mutations using pyrosequencing.

### Image pre-processing

For the Moffitt cohort, H&E FFPE tissue slides were scanned using the Aperio AT2 high-volume digital whole slide scanner at 20× magnification, corresponding to a resolution of 0.5 mu-m pixel-1. For the TCGA cohort, the H&E FFPE tissue slides were downloaded in SVS format from the GDC. Each whole slide image (WSI) was divided into 512-pixel × 512-pixel tiles with no overlap between adjacent tiles. Tiles with more than 50% background pixels were removed (background pixel is defined as a pixel with a gray-scale pixel value greater than 220). Macenko stain normalization [45] was applied to each tile image using a reference WSI to correct for differences in the staining process. Tile images with low contrast or empty tissue masks were removed. For each whole slide image (WSI), the tumor area was annotated by our pathologist (DSV) at a 2× to 4× magnification using the Aperio ImageScope pathology slide viewing software. The tile images were assigned a “tumor” or “benign” label if the entire image was inside or outside the tumor-annotated area, respectively. The “tumor” tile images were assigned a “primary” or “metastatic” label if the WSI tissue was derived from the lung or another tissue, respectively.

### Model training and hyperparameter tuning

We trained and tested four different deep convolutional neural networks (CNNs) on tile images for the three histopathological analysis tasks described in Results using PyTorch. The four different architectures – ResNet18, ResNet34, ResNet50 [24], and Shufflenet-V2 [46] – were chosen because of their widespread use in computer vision and their recent applications to histopathology data. All models were pre-trained on the ImageNet dataset [47]. The last classification layer in each model was replaced with a fully connected linear layer with one output node, and the weights for this linear layer were initialized randomly.

The training data was augmented using random horizontal and vertical flips. Training tile images were sampled inversely proportional to the frequency of their labels to control for imbalance labels in individual tasks. Models were trained with batches of 64 tile images (ResNet50 runs had a mini-batch size of 32 tile images due to memory constraints). Model weights were optimized using binary cross-entropy, the Adam optimizer [48], and a learning rate of 1 × 10e-4, which was the best-performing learning rate on pilot experiments on the molecular profiling task.

For the tumor-normal and primary-met classification tasks, each neural network was trained for three epochs and tested on validation tile images every 200 mini-batches (400 for ResNet50). The final weights for each model were chosen based on the training step that yielded the lowest validation loss. For the molecular classification task, the final weights for each model were chosen based on the training step that yielded the highest slide-level area under the receiver operating curve (AUROC), as was done in prior work on this task [9].

## Cross-validation

To ensure the robustness of model results, cross-validation data folds were created for each classification task. Each model was trained, tuned, and weights were selected using the training and validation data only. All results are reported on the test data averaged across the cross-validation folds.

For the molecular profiling task, we discarded three slides with no tile images inside the tumor-annotated area. We then removed any WSI from a patient that has LCM data; 182 WSIs remain. All WSIs from patients with multiple WSIs (46) or metastatic WSIs (72) were added permanently to the training set. The remaining 85 WSIs consisted of 33 EGFR WSIs and 52 KRAS WSIs. Nineteen KRAS mutated WSIs were chosen randomly and added permanently to the training set in order to balance the dataset. The rest of the WSIs were divided into six different splits of 138 training WSIs, 22 validation WSIs, and 22 test WSIs. The validation and test sets contain an equal number of KRAS mutated and KRAS wild-type WSIs.

To test localization accuracy of models trained for molecular profiling, we augmented the training and test sets of folds 1-5 with 21 WSIs with LCM data (11 with KRAS mutations and 10 without). One of these WSI with a KRAS mutation was randomly chosen and permanently added to the training set. The 20 remaining WSIs were split into five balanced sets, each containing 16 training WSIs and four test WSIs, that were added to folds one through five. All 21 WSIs were added to the training set for fold six.

For the task in which models were trained to identify the primary tissue of a tumor, three WSIs without any tile images inside the tumor-annotated were discarded. Out of the remaining 281 WSIs, 72 WSIs from patients with multiple WSIs were added permanently to the training set. One metastatic WSI and 45 primary lung WSIs were chosen randomly and added permanently to the training set. The remaining 100 WSIs (50 primary and 50 metastatic) were divided into five different splits of 80 training WSIs, 20 validation WSIs, and 20 test WSIs. The validation and test partitions each contain 10 WSIs from primary lung tissue and 10 WSIs from metastatic sites to ensure balanced datasets for evaluation. Like the tumor-normal folds, each of the 100 WSIs was in a validation split and a test split exactly once.

Lastly, for the task in which models were trained to detect tumor, 148 WSIs from primary lung tissue were considered. Any patient with multiple WSIs or at least one WSI with tile images only inside or outside the tumor-annotated area was added permanently to the training set. Of the remaining 93 WSIs, three were randomly chosen to be permanently in the training set. The 90 remaining WSIs were divided into five different splits of 54 training WSIs, 18 validation WSIs, and 18 test WSIs. Each of the 90 WSIs was used in a validation split and a test split exactly once.

## Performance metrics

For each task we calculated weighted accuracy and AUROC, as is standard in recent work in automating histopathological analysis [9]. Weighted accuracy is defined as the average of the true positive rate and true negative rate; a model score of 0.5 is used as the boundary between negative and positive. For the primary-metastatic task and KRAS-mutation task, we also calculated the performance metrics at the slide level. Each WSI is assigned a score based on the median of all the tile-level model scores. All reported performance metrics were computed using the WSIs in the held-out test set. The model scores are concatenated across all cross-validation test folds. If a tile image has multiple model scores because it appears in more than one fold, the median model score is assigned as the final model score.

Confidence intervals for each metric are calculated by bootstrapping (1000 iterations). For an experiment with *N* exemplars in the test set, we sampled N exemplars with replacement. We performed this sampling scheme for 1000 iterations and calculated weighted accuracy and AUROC for each iteration. The distribution of bootstrapped weighted accuracy and AUROC metrics was used to identify the 95% confidence intervals.

## Grad-CAM maps

For each trained model, we visualized the learned analysis strategies using generated guided Grad-CAM [31]. This method generates a feature importance map of the same size as an input image, which indicates the pixels that contributed to the model’s decision for that image. The final convolutional layer in each model (“conv5” for ShuffleNet and “layer4” for the ResNet models) was used to generate the Grad-CAM mask, which ignores noisy locations in the feature importance map. We generated these maps for all tile images in test sets using code from: https://github.com/kazuto1011/grad-cam-pytorch.

Given an input tile image of size 512 × 512 × 3 (three color channels: red, green, blue), the output Grad-CAM map has the same dimensions of 512 × 512 × 3. We processed each Grad-CAM map using the following sequence: (*i*) each value in the map was set to its absolute value, (*ii*) the mean value across feature channels at every location was stored, yielding a 512 × 512 × 1 map, (*iii*) outlier values that were more than three standard deviations away from the mean pixel value in the map were clamped to three standard deviations, and finally *(iv*) the map was normalized to [0,1] for visualization.

## Nuclei segmentation

We used the PyTorch implementation of HoVer-Net to segment and type nuclei in WSIs. The model was trained on the PanNuke dataset [49]. All tile images in the tumor annotated regions of WSIs and all LCM patches were processed using HoVer-Net. The model labels each segmented nuclei as belonging to one of the following categories: unknown, neoplastic, inflammatory, connective, dead, or non-neoplastic epithelial. We combined the “dead” category with the “inflammatory” category based on observations by expert pathologists that the model was misclassifying lymphocytes as “dead”.

**Table 3.**
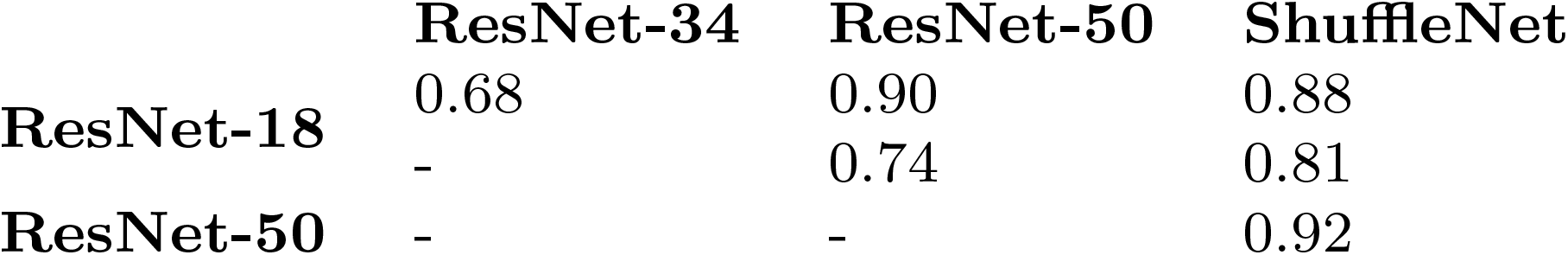
Model decisions are correlated for the KRAS detection task. Pearson’s r correlation coefficients are reported for every model pair tested. All correlation coefficients are statistically significant (*p* < 0.001).

## Nuclei-gradient score calculation

The nuclei-gradient score is based on the intersection over union (IoU) metric that is used to assess object segmentation accuracy in computer vision [50]. For a tile image, we used its Grad-CAM map and the nuclei segmentation results to calculate a nuclei-gradient score for each type of nuclei. Given a threshold value between zero and one, all pixels with Grad-CAM map values greater than or equal to the threshold value were identified. The threshold value was set at 0.5 for results reported in the main text. To compute the IoU, we first counted the suprathreshold Grad-CAM map values. This count was used as the denominator in the IoU. Next, we counted the suprathreshold Grad-CAM map values that also fell within a segmented nucleus, which was used as the numerator in the IoU.

## Supplementary information

### Model misclassifications on tumor detection indicate a reliable visual strategy

We examined 50 image patches in the Moffitt dataset that were misclassified in the tumor detection task. Upon closer examination by expert pathologists, model predictions were in fact correct, and the misclassification reflected imprecise tumor boundaries in the annotations. Indeed, the ground-truth labels on this task did not have cell-level precision. However, the fact that the models could overcome such label noise suggests that the task promotes robust visual strategies.

## Acknowledgments

This study was supported by NIGMS/Advance-CTR through IDeA-CTR grant U54GM115677. Additional support was provided by the Center for Computation and Visualization at Brown (computing hardware supported by NIH Office of the Director grant S10OD025181) and both a State of Florida Endowed Chair in Cancer Research and the DeBartolo Family Personalized Medicine Institute at Moffitt Cancer Center.

---

**Fig. 6 Figure S1:**
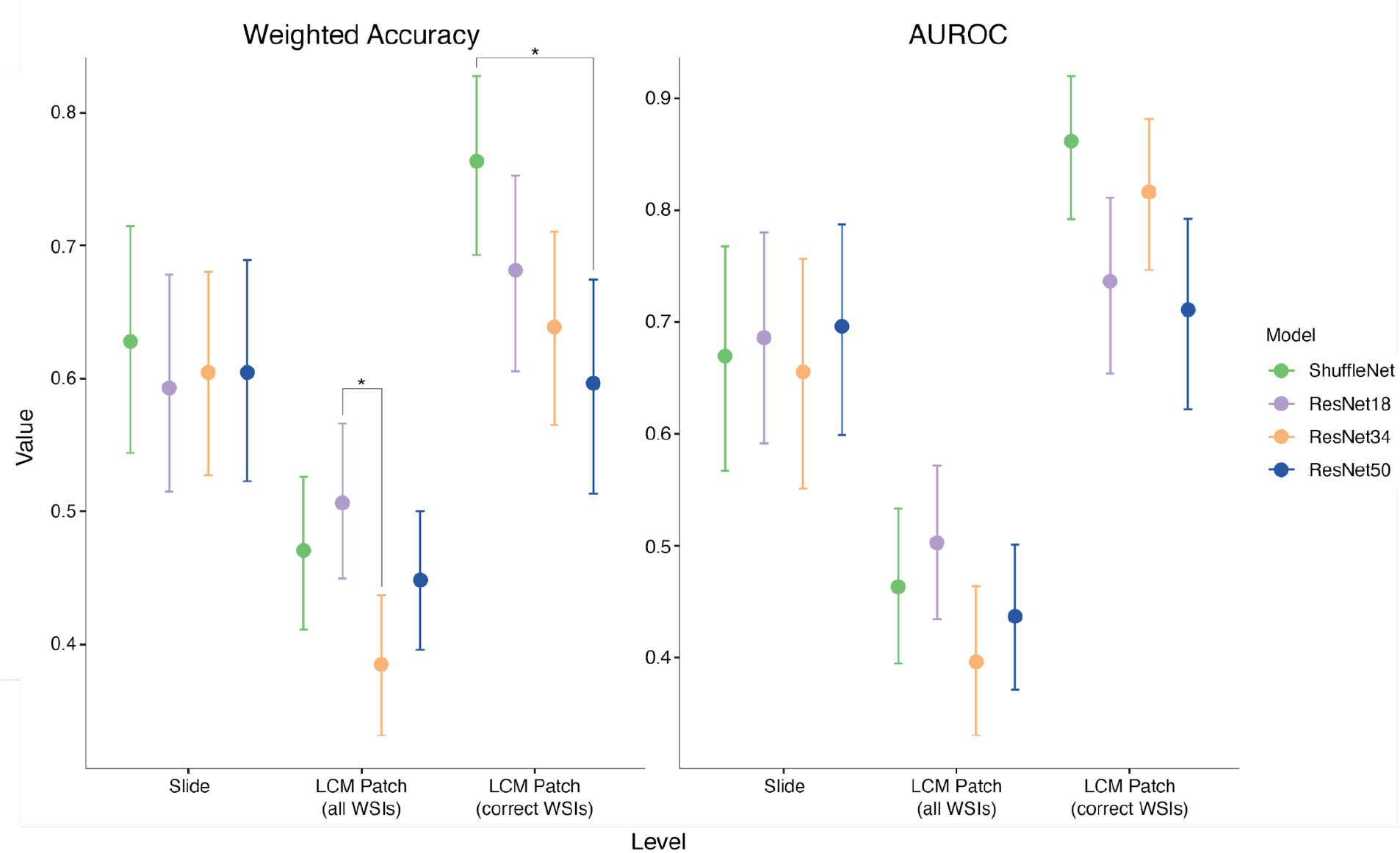
Summary of performance at the slide and patch level for detecting KRAS mutations in lung adenocarcinoma. Point plots showing the performance metrics of four deep learning models at the slide-level and LCM patch-level. The level of the performance metrics is shown on the x-axis, and the value of the metric is on the y-axis. Each point is colored according to the model. The error bars represent the 95% confidence intervals using bootstrapping.

The results published here are in part based upon data generated by the TCGA Research Network: https://www.cancer.gov/tcga.

## References

[1] Rubin, R., Strayer, D.S., Rubin, E., et al.: Rubin’s pathology: clinico-pathologic foundations of medicine (2008)

[2] Esteva, A., Kuprel, B., Novoa, R.A., Ko, J., Swetter, S.M., Blau, H.M., Thrun, S.: Dermatologist-level classification of skin cancer with deep neural networks. Nature 542(7639), 115–118 (2017)

[3] Tschandl, P., Rinner, C., Apalla, Z., Argenziano, G., Codella, N., Halpern, A., Janda, M., Lallas, A., Longo, C., Malvehy, J., Paoli, J., Puig, S., Rosendahl, C., Soyer, H.P., Zalaudek, I., Kittler, H.: Human-computer collaboration for skin cancer recognition. Nat. Med. 26(8), 1229–1234 (2020)

[4] Arcadu, F., Benmansour, F., Maunz, A., Willis, J., Haskova, Z., Prunotto, M.: Deep learning algorithm predicts diabetic retinopathy progression in individual patients. NPJ Digit Med 2, 92 (2019)

[5] Gulshan, V., Peng, L., Coram, M., Stumpe, M.C., Wu, D., Narayanaswamy, A., Venugopalan, S., Widner, K., Madams, T., Cuadros, J., Kim, R., Raman, R., Nelson, P.C., Mega, J.L., Webster, D.R.: Development and validation of a deep learning algorithm for detection of diabetic retinopathy in retinal fundus photographs. JAMA 316(22), 2402–2410 (2016)

[6] Ting, D.S.W., Cheung, C.Y.-L., Lim, G., Tan, G.S.W., Quang, N.D., Gan, A., Hamzah, H., Garcia-Franco, R., San Yeo, I.Y., Lee, S.Y., Wong, E.Y.M., Sabanayagam, C., Baskaran, M., Ibrahim, F., Tan, N.C., Finkelstein, E.A., Lamoureux, E.L., Wong, I.Y., Bressler, N.M., Sivaprasad, S., Varma, R., Jonas, J.B., He, M.G., Cheng, C.-Y., Cheung, G.C.M., Aung, T., Hsu, W., Lee, M.L., Wong, T.Y.: Development and validation of a deep learning system for diabetic retinopathy and related eye diseases using retinal images from multiethnic populations with diabetes. JAMA 318(22), 2211–2223 (2017)

[7] Lotter, W., Diab, A.R., Haslam, B., Kim, J.G., Grisot, G., Wu, E., Wu, K., Onieva, J.O., Boyer, Y., Boxerman, J.L., Wang, M., Bandler, M., Vijayaraghavan, G.R., Gregory Sorensen, A.: Robust breast cancer detection in mammography and digital breast tomosynthesis using an annotation-efficient deep learning approach. Nat. Med. 27(2), 244–249 (2021)

[8] McKinney, S.M., Sieniek, M., Godbole, V., Godwin, J., Antropova, N., Ashrafian, H., Back, T., Chesus, M., Corrado, G.S., Darzi, A., Etemadi, M., Garcia-Vicente, F., Gilbert, F.J., Halling-Brown, M., Hassabis, D., Jansen, S., Karthikesalingam, A., Kelly, C.J., King, D., Ledsam, J.R., Melnick, D., Mostofi, H., Peng, L., Reicher, J.J., Romera-Paredes, B., Sidebottom, R., Suleyman, M., Tse, D., Young, K.C., De Fauw, J., Shetty, S.: International evaluation of an AI system for breast cancer screening. Nature 577(7788), 89–94 (2020)

[9] Coudray, N., Ocampo, P.S., Sakellaropoulos, T., Narula, N., Snuderl, M., Fenyö, D., Moreira, A.L., Razavian, N., Tsirigos, A.: Classification and mutation prediction from non-small cell lung cancer histopathology images using deep learning. Nat. Med. 24(10), 1559–1567 (2018)

[10] Coudray, N., Tsirigos, A.: Deep learning links histology, molecular signatures and prognosis in cancer. Nature Cancer 1(8), 755–757 (2020)

[11] Kather, J.N., Heij, L.R., Grabsch, H.I., Loeffler, C., Echle, A., Muti, H.S., Krause, J., Niehues, J.M., Sommer, K.A.J., Bankhead, P., Kooreman, L.F.S., Schulte, J.J., Cipriani, N.A., Buelow, R.D., Boor, P., Ortiz-Bruchle, N.-N., Hanby, A.M., Speirs, V., Kochanny, S., Patnaik, A., Srisuwananukorn, A., Brenner, H., Hoffmeister, M., van den Brandt, P.A., Jäger, D., Trautwein, C., Pearson, A.T., Luedde, T.: Pan-cancer imagebased detection of clinically actionable genetic alterations. Nat Cancer 1(8), 789–799 (2020)

[12] Fu, Y., Jung, A.W., Torne, R.V., Gonzalez, S., Vöhringer, H., Shmatko, A., Yates, L.R., Jimenez-Linan, M., Moore, L., Gerstung, M.: Pan-cancer computational histopathology reveals mutations, tumor composition and prognosis. Nature Cancer 1(8), 800–810 (2020)

[13] Lu, M.Y., Chen, T.Y., Williamson, D.F.K., Zhao, M., Shady, M., Lipkova, J., Mahmood, F.: AI-based pathology predicts origins for cancers of unknown primary. Nature 594(7861), 106–110 (2021)

[14] Jiao, W., Atwal, G., Polak, P., Karlic, R., Cuppen, E., PCAWG Tumor Subtypes and Clinical Translation Working Group, Danyi, A., de Ridder, J., van Herpen, C., Lolkema, M.P., Steeghs, N., Getz, G., Morris, Q., Stein, L.D., PCAWG Consortium: A deep learning system accurately classifies primary and metastatic cancers using passenger mutation patterns. Nat. Commun. 11(1), 728 (2020)

[15] Geirhos, R., Jacobsen, J.-H., Michaelis, C., Zemel, R., Brendel, W., Bethge, M., Wichmann, F.A.: Shortcut learning in deep neural networks. Nature Machine Intelligence 2(11), 665–673 (2020)

[16] Lapuschkin, S., Wäldchen, S., Binder, A., Montavon, G., Samek, W., Muäller, K.-R.: Unmasking clever hans predictors and assessing what machines really learn. Nat. Commun. 10(1), 1096 (2019)

[17] Pfungst, Oskar., Rahn, Leo, C.: Clever Hans (the Horse of Mr. Von Osten.) a Contribution to Experimental Animal and Human Psychology,. Holt, Rinehart and Winston, New York (1911)

[18] Diao, J.A., Wang, J.K., Chui, W.F., Mountain, V., Gullapally, S.C., Srinivasan, R., Mitchell, R.N., Glass, B., Hoffman, S., Rao, S.K., Maheshwari, C., Lahiri, A., Prakash, A., McLoughlin, R., Kerner, J.K., Resnick, M.B., Montalto, M.C., Khosla, A., Wapinski, I.N., Beck, A.H., Elliott, H.L., Taylor-Weiner, A.: Human-interpretable image features derived from densely mapped cancer pathology slides predict diverse molecular phenotypes. Nat. Commun. 12(1), 1–15 (2021)

[19] Mobadersany, P., Yousefi, S., Amgad, M., Gutman, D.A., Barnholtz-Sloan, J.S., Velázquez Vega, J.E., Brat, D.J., Cooper, L.A.D.: Predicting cancer outcomes from histology and genomics using convolutional networks. Proc. Natl. Acad. Sci. U. S. A. 115(13), 2970–2979 (2018)

[20] Saltz, J., Gupta, R., Hou, L., Kurc, T., Singh, P., Nguyen, V., Samaras, D., Shroyer, K.R., Zhao, T., Batiste, R., Van Arnam, J., Cancer Genome Atlas Research Network, Shmulevich, I., Rao, A.U.K., Lazar, A.J., Sharma, A., Thorsson, V.: Spatial organization and molecular correlation of Tumor-Infiltrating lymphocytes using deep learning on pathology images. Cell Rep. 23(1), 181–1937 (2018)

[21] Rekhtman, N., Ang, D.C., Riely, G.J., Ladanyi, M., Moreira, A.L.: KRAS mutations are associated with solid growth pattern and tumor-infiltrating leukocytes in lung adenocarcinoma. Mod. Pathol. 26(10), 1307–1319 (2013)

[22] Kather, J.N., Pearson, A.T., Halama, N., Jäger, D., Krause, J., Loosen, S.H., Marx, A., Boor, P., Tacke, F., Neumann, U.P., Grabsch, H.I., Yoshikawa, T., Brenner, H., Chang-Claude, J., Hoffmeister, M., Trautwein, C., Luedde, T.: Deep learning can predict microsatellite instability directly from histology in gastrointestinal cancer. Nat. Med. 25(7), 1054–1056 (2019)

[23] Espina, V., Wulfkuhle, J.D., Calvert, V.S., VanMeter, A., Zhou, W., Coukos, G., Geho, D.H., Petricoin, E.F., Liotta, L.A.: Laser-capture microdissection. Nat. Protoc. 1(2), 586–603 (2006)

[24] He, K., Zhang, X., Ren, S., Sun, J.: Deep residual learning for image recognition (2015) https://arxiv.org/abs/1512.03385 [cs.CV]

[25] Boyle, T.A., Mondal, A.K., Saeed-Vafa, D., Ananth, S., Ahluwalia, P., Kothapalli, R., Chaubey, A., Roberts, E., Qin, D., Magliocco, A.M., Rojiani, A.M., Kolhe, R.: Guideline-Adherent clinical validation of a comprehensive 170-gene DNA/RNA panel for determination of small variants, copy number variations, splice variants, and fusions on a Next-Generation sequencing platform in the CLIA setting. Front. Genet. 12, 503830 (2021)

[26] Linsley, D., Eberhardt, S., Sharma, T., Gupta, P., Serre, T.: What are the visual features underlying human versus machine vision? In: 2017 IEEE International Conference on Computer Vision Workshops (ICCVW), pp. 2706–2714 (2017)

[27] Zeiler, M.D., Fergus, R.: Visualizing and understanding convolutional networks (2013) https://arxiv.org/abs/1311.2901 [cs.CV]

[28] Linsley, D., Shiebler, D., Eberhardt, S., Serre, T.: Learning what and where to attend (2019)

[29] Simonyan, K., Vedaldi, A., Zisserman, A.: Deep inside convolutional networks: Visualising image classification models and saliency maps (2013) https://arxiv.org/abs/1312.6034 [cs.CV]

[30] Linsley, J.W., Linsley, D.A., Lamstein, J., Ryan, G., Shah, K., Castello, N.A., Oza, V., Kalra, J., Wang, S., Tokuno, Z., Javaherian, A., Serre, T., Finkbeiner, S.: Superhuman cell death detection with biomarker-optimized neural networks. Sci Adv 7(50), 8142 (2021)

[31] Selvaraju, R.R., Cogswell, M., Das, A., Vedantam, R., Parikh, D., Batra, D.: Grad-CAM: Visual explanations from deep networks via Gradient-Based localization. In: 2017 IEEE International Conference on Computer Vision (ICCV), pp. 618–626 (2017)

[32] Graham, S., Vu, Q.D., Raza, S.E.A., Azam, A., Tsang, Y.W., Kwak, J.T., Rajpoot, N.: Hover-Net: Simultaneous segmentation and classification of nuclei in multi-tissue histology images. Med. Image Anal. 58, 101563 (2019)

[33] Kumar, V., Abbas, A.K., Aster, J.C.: Robbins Basic Pathology E-Book. Elsevier Health Sciences, ??? (2017)

[34] Noorbakhsh, J., Farahmand, S., Foroughi Pour, A., Namburi, S., Caruana, D., Rimm, D., Soltanieh-Ha, M., Zarringhalam, K., Chuang, J.H.: Deep learning-based cross-classifications reveal conserved spatial behaviors within tumor histological images. Nat. Commun. 11(1), 6367 (2020)

[35] Edgington, E.S.: RANDOMIZATION TESTS. J. Psychol. 57, 445–449 (1964)

[36] Wei, J.W., Tafe, L.J., Linnik, Y.A., Vaickus, L.J., Tomita, N., Hassanpour, S.: Pathologist-level classification of histologic patterns on resected lung adenocarcinoma slides with deep neural networks. Sci. Rep. 9(1), 3358 (2019)

[37] Gertych, A., Swiderska-Chadaj, Z., Ma, Z., Ing, N., Markiewicz, T., Cierniak, S., Salemi, H., Guzman, S., Walts, A.E., Knudsen, B.S.: Convolutional neural networks can accurately distinguish four histologic growth patterns of lung adenocarcinoma in digital slides. Sci. Rep. 9(1), 1483 (2019)

[38] Travis, W.D., Brambilla, E., Noguchi, M., Nicholson, A.G., Geisinger, K., Yatabe, Y., Ishikawa, Y., Wistuba, I., Flieder, D.B., Franklin, W., Gazdar, A., Hasleton, P.S., Henderson, D.W., Kerr, K.M., Petersen, I., Roggli, V., Thunnissen, E., Tsao, M.: Diagnosis of lung cancer in small biopsies and cytology: implications of the 2011 international association for the study of lung Cancer/American thoracic Society/European respiratory society classification. Arch. Pathol. Lab. Med. 137(5), 668–684 (2013)

[39] Campanella, G., Hanna, M.G., Geneslaw, L., Miraflor, A., Silva, V.W.K., Busam, K.J., Brogi, E., Reuter, V.E., Klimstra, D.S., Fuchs, T.J.: Clinical-grade computational pathology using weakly supervised deep learning on whole slide images. Nat. Med. 25(8), 1301–1309 (2019)

[40] Lee, K., Zung, J., Li, P., Jain, V., Sebastian Seung, H.: Superhuman accuracy on the SNEMI3D connectomics challenge (2017) https://arxiv.org/abs/1706.00120 [cs.CV]

[41] DeGrave, A.J., Janizek, J., Lee, S.-I.: AI for radiographic COVID-19 detection selects shortcuts over signal. Nature Machine Intelligence 3(7), 610–619 (2021)

[42] Linsley, D., Kim, J., Ashok, A., Serre, T.: Recurrent neural circuits for contour detection. International Conference on Learning Representations (2020)

[43] Knepper, T.C., Bell, G.C., Hicks, J.K., Padron, E., Teer, J.K., Vo, T.T., Gillis, N.K., Mason, N.T., McLeod, H.L., Walko, C.M.: Key lessons learned from moffitt’s molecular tumor board: The clinical genomics action committee experience. Oncologist 22(2), 144–151 (2017)

[44] Ellrott, K., Bailey, M.H., Saksena, G., Covington, K.R., Kandoth, C., Stewart, C., Hess, J., Ma, S., Chiotti, K.E., McLellan, M., Sofia, H.J., Hutter, C., Getz, G., Wheeler, D., Ding, L., MC3 Working Group, Cancer Genome Atlas Research Network: Scalable open science approach for mutation calling of tumor exomes using multiple genomic pipelines. Cell Syst 6(3), 271–2817 (2018)

[45] Macenko, M., Niethammer, M., Marron, J.S., Borland, D., Woosley, J.T., Guan, X., Schmitt, C., Thomas, N.E.: A method for normalizing histology slides for quantitative analysis. In: 2009 IEEE International Symposium on Biomedical Imaging: From Nano to Macro, pp. 1107–1110 (2009)

[46] Ma, N., Zhang, X., Zheng, H.-T., Sun, J.: ShuffleNet v2: Practical guidelines for efficient CNN architecture design (2018) https://arxiv.org/abs/1807.11164 [cs.CV]

[47] Deng, J., Dong, W., Socher, R., Li, L.-J., Li, K., Fei-Fei, L.: ImageNet: A large-scale hierarchical image database. In: 2009 IEEE Conference on Computer Vision and Pattern Recognition, pp. 248–255 (2009)

[48] Kingma, D.P., Ba, J.: Adam: A method for stochastic optimization. arXiv preprint arXiv:1412.6980 (2014)

[49] Gamper, J., Koohbanani, N.A., Benes, K., Graham, S., Jahanifar, M., Khurram, S.A., Azam, A., Hewitt, K., Rajpoot, N.: PanNuke dataset extension, insights and baselines (2020) https://arxiv.org/abs/2003.10778 [eess.IV]

[50] Lin, T.-Y., Maire, M., Belongie, S., Bourdev, L., Girshick, R., Hays, J., Perona, P., Ramanan, D., Lawrence Zitnick, C., Dollár, P.: Microsoft COCO: Common objects in context (2014) https://arxiv.org/abs/1405.0312 [cs.CV]

